# qPCR Guru, a free browser-based platform, strengthens microRNA analyses using full-curve Cq estimation

**DOI:** 10.64898/2026.06.21.733609

**Authors:** Amar M. Singh, Owen Singh, MD Marzan Sarkar, Rob Coultous, Steven L. Stice

## Abstract

Quantitative PCR (qPCR) depends on reliable quantification cycle (Cq) estimation from amplification curves, which are not always well-behaved. We developed qPCR Guru to provide a complete analysis pipeline including data quality assessment, relative quantification, standard-curve diagnostics, and dual-method Cq evaluation. The latter compares the conventional instrument-derived threshold (“Reported”) Cq versus the full-curve five-parameter logistic (5PL) second-derivative-maximum (“Fit”) Cq and automatically flags curve-shape abnormalities and disagreement between the two estimates. On high-expressing targets (mRNA and microRNA), the two methods showed strong convergence, confirming general-purpose performance. On low-expressing targets, such as serum microRNA, baseline artifacts and biphasic amplification result in threshold miscalls that standard instrument analysis does not flag. Fit Cq restored replicate-concordant values where Reported Cq split the technical replicates by 17–20 cycles, recovered MIQE-compliant amplification efficiencies lost to biphasic miscalls (from 74% to 102% and 387% to 98%), and lowered within-group variability by 48% and 68% in feline and bovine samples, respectively. Together, these results demonstrate that full-curve estimation, with integrated curve-level diagnostics, strengthens qPCR analyses against threshold miscalls.

**ARTICLE HIGHLIGHTS:**

- qPCR Guru is a free, browser-based platform that provides a complete analysis pipeline and facilitates side-by-side comparisons of an instrument’s threshold (Reported) Cq and a full-curve (Fit) Cq, from the five-parameter logistic fitting with second-derivative-maximum (SDM/cpD2).
- For every well the application automatically flags curve-shape abnormalities and disagreement between the two Cq estimates.
- On clean, high-expressing mRNA and microRNA targets, the two estimators (Reported Cq and Fit Cq) were strongly concordant and produced equivalent relative quantification with comparable precision.
- In low-expressing serum microRNA, baseline artifacts and biphasic amplification produced threshold Cq miscalls of up to ∼20 cycles and were detected by curve-shape flags and/or method disagreement.
- The full-curve Cq estimate recovered replicate-concordant values, restored MIQE-compliant amplification efficiencies, and reduced within-group variability in serum microRNA.

## INTRODUCTION

Quantitative polymerase chain reaction (qPCR) is the gold-standard method of nucleic acid quantification, valued for its sensitivity, dynamic range, and ease of use [1, 2]. Among its applications, the quantification of microRNAs (miRNAs), short ∼22-nucleotide regulators of gene expression that circulate stably in serum and other biofluids, has drawn widespread interest, both for the study of post-transcriptional regulation and as a source of minimally invasive biomarkers for diagnostic and prognostic clinical applications [3, 4]. Quantifying miRNAs by reverse transcription qPCR (RT-qPCR) poses significant challenges not typically encountered with messenger RNAs (mRNA) due to size, normalization, and abundance. The reverse-transcription methodologies used to convert miRNAs into complementary DNA (cDNA) templates, such as the stem-loop and polyadenylation-based approaches [5, 6], result in short amplicons that, when coupled with the low template abundance common in biofluids, frequently yield amplification curves that are not as well-behaved as those from abundant, longer targets.

Irrespective of the target, determining the concentration of a DNA or RNA molecule by qPCR depends on the successful identification of a quantification cycle (Cq) using its amplification curve, from which the relative or absolute levels of the target can be derived [7, 8]. The Cq is conventionally determined as the point at which fluorescence crosses a fixed threshold within the exponential phase, an approach that performs reliably when the curve has a stable baseline and a well-defined exponential rise. Full-curve approaches, such as the five-parameter logistic (5PL) method, fit a sigmoidal curve to the fluorescence amplification values and derive the Cq from the fitted curve’s second-derivative maximum (SDM or cpD2), and have demonstrated advantages for qPCR data modeling [9, 10]. Such methods remain comparatively underused, as threshold values are produced automatically by instrument software whereas full-curve analysis has typically required specialized tools and access to the raw fluorescence data [11, 12]. Reliable estimation matters beyond the individual value: amplification efficiency and replicate precision, both required for credible reporting under the MIQE guidelines [13], are themselves derived from Cq, yet adherence to such reporting standards remains limited [14].

While full-curve estimation is established for qPCR broadly, its specific consequences for the low-input, artifact-prone amplification encountered in miRNA assays have not been defined. Here, we describe qPCR Guru, a free, browser-based platform that places two Cq estimates side by side for every well: the conventional instrument-derived threshold value (referred to as the “Reported Cq”) and a full-curve 5PL second-derivative-maximum fit value, referred to as the “Fit Cq (SDM)”. The platform additionally flags curve-shape abnormalities and disagreement between the two estimates. Using this dual-method comparison, we evaluate where the two approaches agree and where they diverge across high- and low-expressing targets spanning mRNA and serum miRNA.

## METHODS

### Samples, RNA isolation and RT-qPCR

Datasets from previous RT-qPCR studies evaluating Akt knockdown in CT-2A mouse glioma cells were made available for analysis [15]. Feline serum was purchased from BioIVT (Woodbury, NY), using individual samples or a pooled source of 6 animals. Bovine serum, pooled source, was purchased from Fisher Scientific (Pittsburgh, PA). MicroRNA were extracted using the miRNeasy serum/plasma advanced kit (Qiagen, Germantown, MD) according to manufacturer instructions. Reverse transcription was performed using the TaqMan Advanced MicroRNA cDNA Synthesis Kit with poly-A tailing and RNA adapter ligation, according to manufacturer’s instructions. All qPCR assays were performed on a QuantStudio 5 (Applied Biosystems, Waltham, MA, USA). TaqMan assays were purchased against microRNA targets for *miR-126*, *miR-28*, *miR-1*, *miR-23a*; *cel-miR-39* co-diluted spike-in control (Fisher Scientific, Pittsburgh, PA). Raw per-cycle fluorescence and instrument-reported Cq values were exported as XLS/XLSX, TXT and RDML for analysis in qPCR Guru (https://qpcrguru.com).

### Dual quantification-cycle estimation

For every well, two quantification-cycle (Cq) values were obtained. The Reported Cq is the threshold-crossing value computed by the real-time PCR instrument software, used as exported without modification. The Fit Cq was derived independently by full-curve modeling, as follows.

Background-subtracted fluorescence F as a function of cycle x was fitted with the five-parameter log-logistic (5PL) model

F(x) = c + (d − c) / [1 + exp(b(ln x − ln e))]^f

where c and d are the lower (baseline) and upper (plateau) asymptotes, b is the Hill slope, e is the inflection-related cycle (equal to the inflection point only when f = 1), and f is an asymmetry parameter [9, 10]. Parameters were estimated by Levenberg–Marquardt nonlinear least squares, with the asymmetry parameter constrained to a smooth bounded range to prevent degenerate solutions while leaving derived metrics unaffected. Goodness of fit was recorded as the coefficient of determination, R² (5PL). The Fit Cq was defined as the second-derivative maximum (SDM, equivalent to qpcR’s cpD2) of the fitted curve, located numerically from a five-point central-difference estimate of F″(x); the SDM was used in preference to the first-derivative maximum because it agrees with threshold-based Cq to within approximately one cycle for well-behaved curves [9]. For each well, the difference between estimators was recorded as ΔCq = Fit Cq − Reported Cq.

The 5PL implementation was validated against the qpcR R package [9]; fitted parameters and derived Cq values agreed to four decimal places across the test set.

### Curve quality control

Each well received a curve-shape classification (Normal or Abnormal) derived from the fit and the raw trajectory. Abnormal classifications captured curves that failed to converge, showed poor fit, lacked genuine amplification, or were biphasic. Biphasic curves — two distinct amplification events separated by an intervening plateau, characteristic of primer-dimer artifacts or low-template stochastic amplification — were detected directly from the raw fluorescence by slope-peak analysis rather than from the 5PL fit, because a single-sigmoid model can absorb a secondary rise into an elevated baseline while still returning a high R². To avoid inconsistent verdicts between near-identical replicates, each flag required multiple corroborating signals (fit quality, curve shape, and amplitude) rather than a single threshold. Within replicate groups of sufficient size, additional outlier wells were identified from the dispersion of fitted inflection cycles using a median-absolute-deviation criterion floored at 0.5 cycles.

Separately from curve shape, wells were flagged for method disagreement when |ΔCq| met a fixed threshold, identifying threshold miscalls not captured by shape classification alone.

### Relative quantification and amplification efficiency

qPCR Guru computes both Cq estimates for every well and can perform relative quantification using either, by the comparative-Cq method [7] with optional efficiency correction [8]. The ΔΔCq results presented here were generated using the Reported Cq, with the Fit Cq computed in parallel for comparison. Amplification efficiency was determined from standard-curve dilution series by regressing Cq on log₁₀ template dilution and computing E = 10^(−1/slope) − 1, expressed as a percentage [13]; efficiency was calculated separately from Reported and Fit Cq to compare estimators. The *cel-miR-39* was co-diluted across the series as a dilution-series control rather than a constant-input reference.

### Statistical analysis

Within-group precision was quantified as the standard deviation (SD) of Cq across replicates within each sample × target group, computed separately for Reported and Fit Cq. The two estimators were compared by the Wilcoxon signed-rank test on the paired per-group differences (Reported SD − Fit SD), with the reduction in median SD reported as a percentage. Method concordance on clean (high-expressing) data was assessed by Pearson (r) and Spearman (ρ) correlation, Bland–Altman bias and limits of agreement, and Lin’s concordance correlation coefficient for ΔΔCq. All tests were two-sided with significance at α = 0.05. Numeric results are reported as absolute values alongside derived percentages. Analyses were performed in qPCR Guru (version 1.0) for primary analysis along with R (version 4.6.0) and qpcR for cross-validation.

## RESULTS AND DISCUSSION

### Establishment of a dual Cq-estimator analysis platform

qPCR Guru is a browser-based application that enables direct uploads from instrument export files (XLS/XLSX, RDML, CSV, TXT), performs data-quality and melt-peak assessment, fits each amplification curve with the 5PL model, and reports a paired Reported Cq and Fit Cq for every well alongside relative (ΔΔCq) or absolute quantification, standard-curve diagnostics, figure export, and MIQE publication-readiness checks (Figure 1A). The application is designed to capture the instrument-derived Reported Cq, while also deriving a Fit Cq (SDM) from the uploaded data. Depending on whether the curve has a traditional well-behaved amplification (Figure 1B) or an atypical curve such as biphasic amplification (Figure 1C), in which the threshold is crossed by an early non-specific shoulder before the main exponential rise, the two Cq values may converge or diverge. The application reports these data, so a user may quickly and easily compare values, and evaluate curve shapes and flags summarized for each run (Figure 1D).

**Figure 1.**
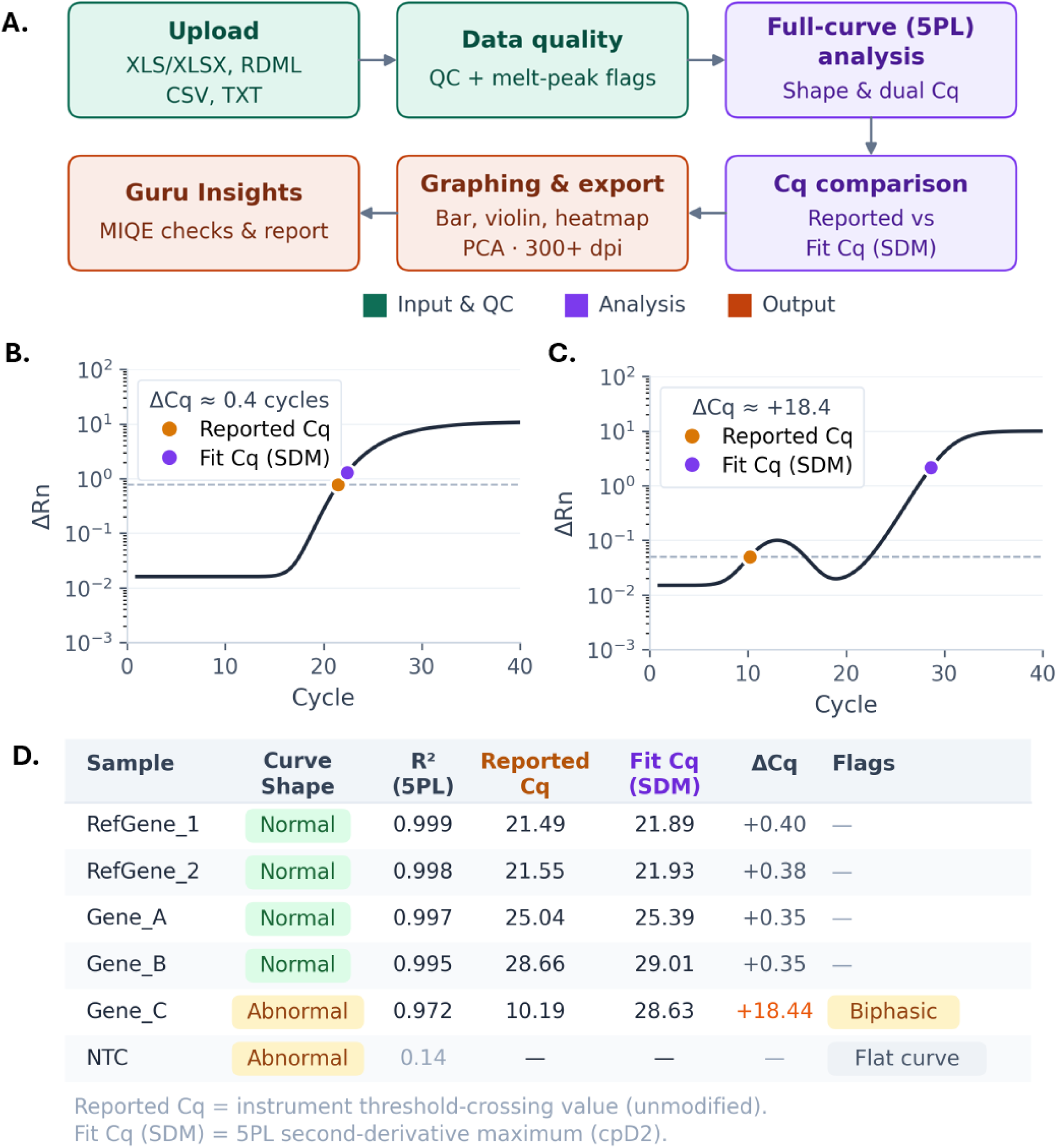
The qPCR Guru platform and dual quantification-cycle estimation. (A) Schematic of the analysis workflow: (1) instrument files (XLS/XLSX, RDML, CSV, TXT) are imported and undergo data-quality and melt-peak assessment, (2) a 5PL curve fitting with curve-shape classification and dual Cq estimation is performed for comparison, and (3) relative quantification, graphing, figure export, MIQE publication-readiness checks and a data assessment, termed Guru Insights, is made available. (B) Illustration of a well-behaved curve, the instrument threshold (“Reported”) Cq (orange) and the full-curve five-parameter logistic second-derivative-maximum (“Fit”) Cq (purple) nearly coincide. (C) Illustration of an abnormal, biphasic curve, where the threshold is crossed by an early shoulder well before the main exponential rise, so the two estimates diverge. (D) Example per-well summary reporting curve-shape classification, 5PL goodness of fit (R²), Reported Cq, Fit Cq (SDM), their difference (ΔCq), and QC flags. Cq, quantification cycle; 5PL, five-parameter logistic; SDM, second-derivative maximum; ΔRn, baseline-corrected normalized fluorescence; NTC, no-template control.

### Full-curve estimation reproduces conventional quantification on well-behaved targets

We initially assessed general-purpose performance using a previous dataset that tested an Akt3 siRNA-knockdown in mouse CT-2A glioma cells across a panel of eight genes (Supplementary Table 1) [15]. Data-quality assessment passed for all targets except *Fasl*, a low-expressing gene flagged for elevated replicate variability and a control-well outlier (Figure 2A, Supplementary Figure 1). Transcript levels for the target genes from knockdown or control cells were assessed by RT-qPCR by the conventional Reported Cq. As expected, *Akt3* was specifically reduced to approximately 15% of control, while the non-targeted *Akt1* and *Akt2* and the remaining pathway genes remained near control levels (Figure 2B–C). Re-running quantification with the Fit Cq produced concordant results (Supplementary Table 2). Across this clean, high-expressing dataset the two estimates were strongly correlated (Pearson r = 0.986, Spearman ρ = 0.996; Figure 3A), and Bland–Altman analysis showed only a small systematic offset (mean +0.81 cycle; limits of agreement −1.39 to +3.00; Figure 3B), consistent with the tendency of the second-derivative maximum to fall slightly later than the threshold crossing [9]. This offset was largest for *Fasl*, a low-expressing, QC-flagged target (Supplementary Figure 1), but because its change was consistent across replicates the differences were largely cancelled in relative quantification, and fold-change estimates from the two methods were essentially identical (Lin’s concordance correlation coefficient = 0.989; Figure 3C). Within-group precision was comparable between methods on this dataset, with no significant difference (Figure 3D). Overall these data indicate that on well-behaved high-expressing targets the choice of Cq estimator (Reported vs Fit) did not materially change the result, and that full-curve estimation reproduces conventional quantification without distorting the data.

**Figure 2.**
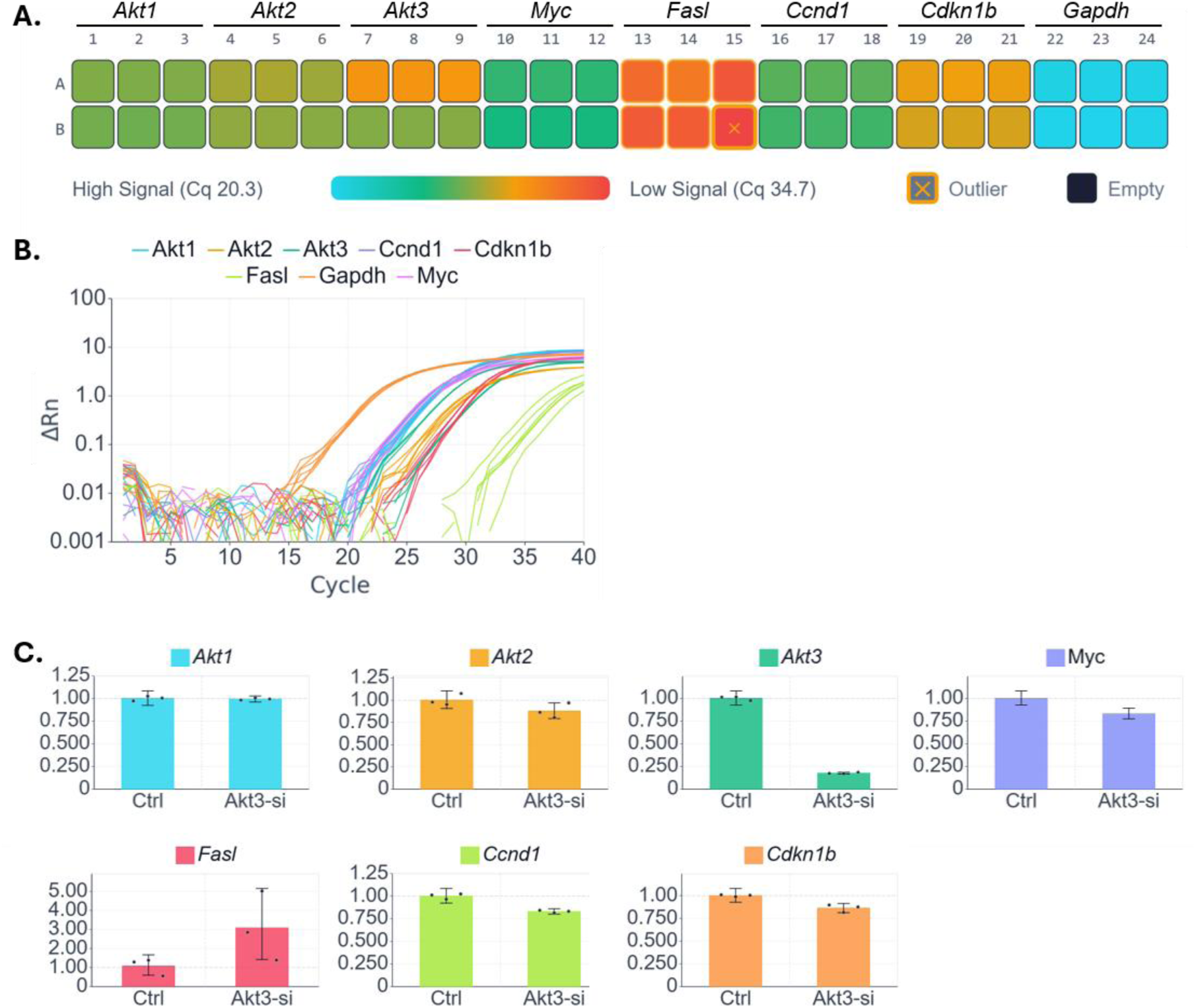
Analysis output from qPCR Guru from an Akt3 knockdown experiment in CT-2A mouse glioma cells. (A) Plate-level quality-control heatmap of Reported Cq across the eight-gene panel (high signal/low Cq, blue; low signal/high Cq, red); the platform-identified outlier well is marked. (B) Amplification curves (ΔRn versus cycle) for all wells, colored by target. (C) Relative quantification (ΔΔCq from Reported Cq) of each gene in Akt3 siRNA-treated versus control cells, normalized to Gapdh; Akt3 is specifically reduced while non-targeted genes remain near control levels. Data is graphed as the mean with standard deviation (n=3, technical replicates).

**Figure 3.**
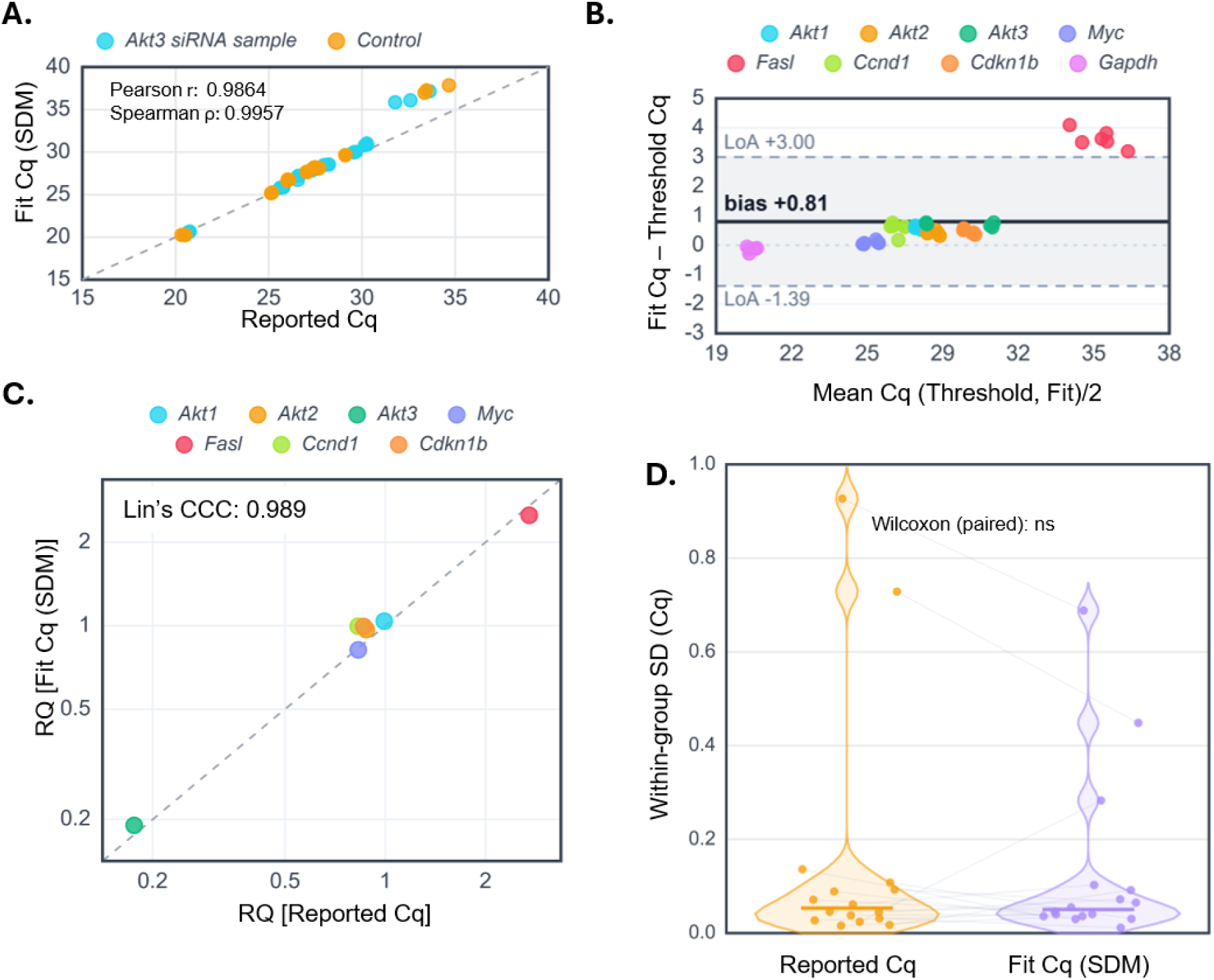
Concordance of the Reported Cq and Fit Cq (SDM) using the Akt3 knockdown dataset. (A) Per-well Fit Cq (SDM) versus Reported Cq for siRNA-treated and control samples show high correlation (Pearson r = 0.986; Spearman ρ = 0.996). (B) Bland–Altman plot of the between-method difference against the mean of the two estimates; solid line, mean bias (+0.81 cycle); dashed lines, 95% limits of agreement (−1.39 to +3.00); the largest offsets correspond to the low-expressing, QC-flagged *Fasl* target. (C) Scatter plot evaluating relative quantification computed from Fit Cq versus from Reported Cq for each gene (Lin’s concordance correlation coefficient = 0.989), indicating high correlation between analysis methods. (D) Bean plots examining within-group standard deviation (SD) of Cq by method; paired Wilcoxon signed-rank test, not significant (ns); connecting lines link the same sample × target group across methods.

### Threshold miscalls occur in low-expressing serum microRNA

During the evaluation of miRNA expression by RT-qPCR in feline serum, we uncovered that unlike the experiments with well-behaved curves on mRNA, the two methods diverged markedly on low-expressing serum miRNA. In well A10 (*miR-126*, sample S1), the instrument threshold was crossed near cycle 12 by an early biphasic shoulder, roughly 18 cycles before the true exponential rise that the Fit Cq tracked (Reported 11.9 vs Fit 30.9; Figure 4A, Table 1, Supplementary Figure 2A,B), whereas its technical replicate twin B10 amplified normally and the two estimates agreed (Figure 4B). A comparable miscall occurred for *miR-28* (sample S4): in well L7 a noisy baseline produced a Reported Cq of 9.9 against a Fit Cq of 30.0, while its technical twin K7 was normal (Figure 4C,D; Table 1, Supplementary Figure 2C,D). These two miscalls differ in an important way. The biphasic A10 curve was flagged as Abnormal by curve-shape classification, but the noisy-baseline L7 curve passed shape classification and was identified only by the large method disagreement (ΔCq) — the case that motivates reporting both estimates rather than relying on curve shape alone. The instrument’s own amplification-confidence scores corroborated both miscalls (A10, 0.101; L7, 0.018; versus approximately 0.42 - 0.46 for their normal twins; Supplementary Figure 2). In each case the Fit Cq returned a value concordant with the well’s normal replicate, recovering values that threshold calling had split by 17–20 cycles. Together, these data demonstrate two instances when the use of a Fit Cq (SDM) may offer advantages over the Reported Cq for data analysis, due to an atypical amplification resulting in threshold miscalls.

**Figure 4.**
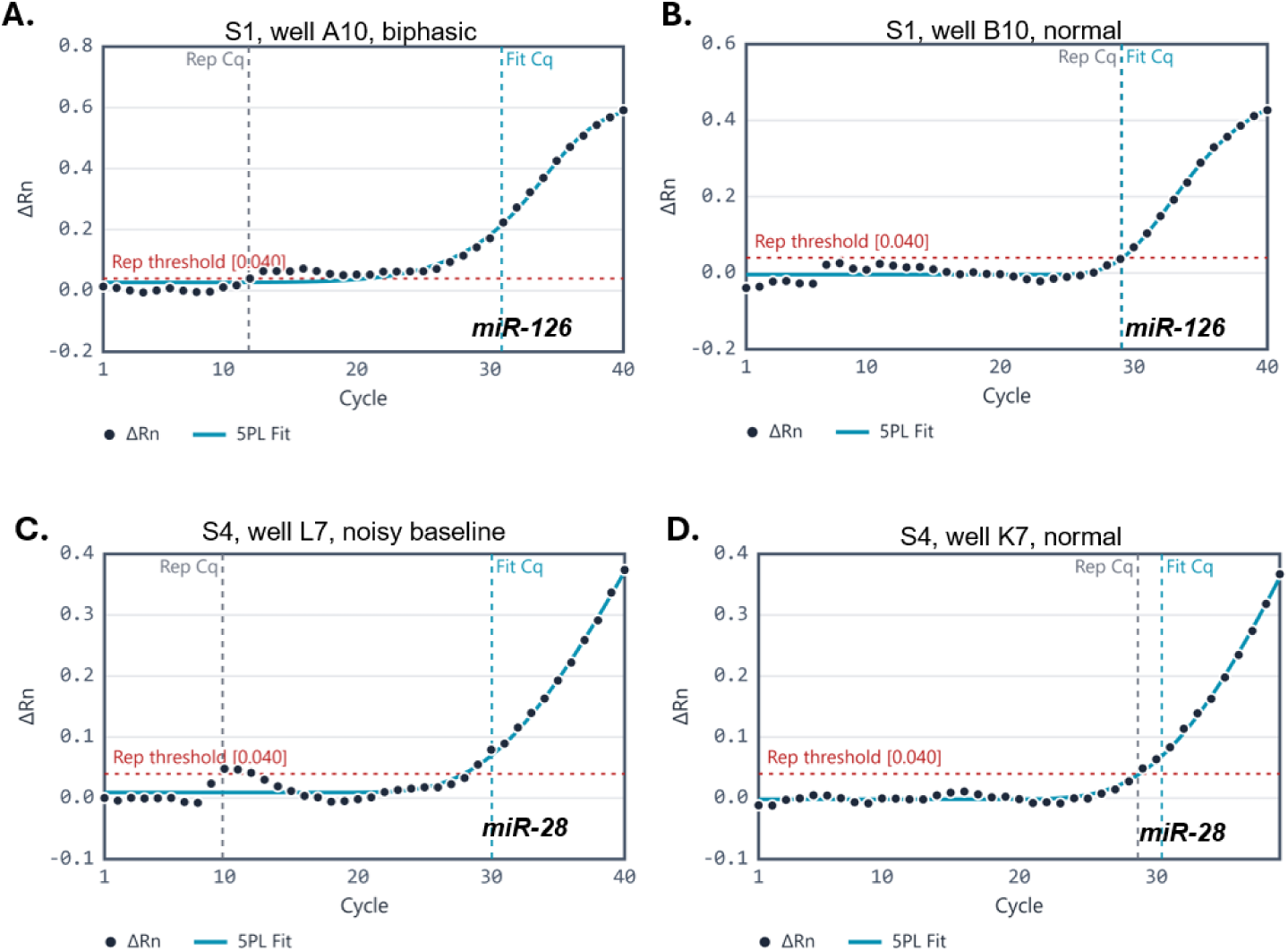
Threshold miscalls in low-expressing serum microRNA recovered by full-curve estimation. Fit curve amplification plots from individual feline serum samples (n=1 per sample) for (A) *miR-126*, sample serum (S) 1, well A10: a biphasic curve in which the threshold is crossed near cycle 12 by an early shoulder, ∼18 cycles before the main rise tracked by the Fit Cq, (B) *miR-126*, sample S1, well B10: the normal technical replicate, for which the two estimates agree, (C) *miR-28*, sample S4, well L7: a noisy-baseline miscall (Reported 9.9 versus Fit 30.0) that passed curve-shape classification, and (D) *miR-28*, sample S4, well K7: the normal technical replicate. Black points, instrument-reported baseline corrected ΔRn per cycle; blue line, 5PL fit; grey dashed line, Reported Cq (instrument threshold; “Rep Cq”); blue dashed line, Fit Cq (SDM); red dotted line, instrument threshold value.

**Table 1.** Dual-Cq comparison for serum miRNA technical replicates. Per-well Reported Cq (instrument threshold) and Fit Cq (SDM) for *miR-126* (samples S1–S2) and *miR-28* (S3–S4) serum replicates. Highlighted wells are threshold miscalls: A10 is flagged as biphasic by 5PL curve-shape QC, whereas L7 carries a Normal curve-shape classification yet miscalls by ∼20 cycles — detectable only by the large ΔCq (Fit − Reported). In both, Fit Cq returns a value concordant with the well’s normal technical twin.^¹^ Curve shape determined by five-parameter logistic (5PL) sigmoid fit (Spiess *et al.* 2008; Ritz & Spiess 2008).^²^ Reported Cq: the instrument threshold-crossing value as imported; unmodified by qPCR Guru.^³^ Fit Cq (SDM): the second-derivative maximum (cpD2) of the fitted 5PL curve (Ritz & Spiess 2008), calculated by qPCR Guru.

| Well | Sample | Target | Curve Shape<sup>1</sup> | R<sup>2</sup> (5PL) | Reported Cq<sup>2</sup> | Fit Cq (SDM)<sup>3</sup> | $\Delta Cq$ | Flags |
| --- | --- | --- | --- | --- | --- | --- | --- | --- |
| A10 | S1 | *miR-126* | Abnormal | 0.988 | 11.91 | 30.88 | +18.97 | Biphasic amplification |
| B10 | S1 | *miR-126* | Normal | 0.988 | 29.05 | 29.11 | +0.06 | — |
| C10 | S2 | *miR-126* | Normal | 0.995 | 29.03 | 29.97 | +0.94 | — |
| D10 | S2 | *miR-126* | Normal | 0.997 | 29.58 | 30.37 | +0.79 | — |
| E7 | S3 | *miR-28* | Normal | 0.982 | 29.21 | 28.76 | -0.45 | — |
| F7 | S3 | *miR-28* | Normal | 0.994 | 29.36 | 28.77 | -0.59 | — |
| K7 | S4 | *miR-28* | Normal | 0.998 | 28.65 | 30.06 | +1.41 | — |
| L7 | S4 | *miR-28* | Normal | 0.983 | 9.85 | 30.04 | +20.19 | — |

### Consequences of threshold miscalls on amplification efficiency

The potential for threshold miscalls to propagate to MIQE-relevant metrics was evaluated. In a standard-curve dilution series, a single biphasic miscall distorted the threshold-based efficiency for *miR-23a*, a miscalled well (D5) inverted the dilution series and yielded an uninterpretable efficiency of 387% (R² = 0.11), whereas the Fit Cq restored a linear standard curve (efficiency 98.4%, R² = 0.93; Figure 5A, Table 2). A similar but less extreme recovery occurred for *miR-1* (74.4% to 102.0%; Figure 5B). The co-diluted *cel-miR-39* control, which amplified cleanly, passed by both methods (103.5% vs 93.5%; Figure 5C), confirming that full-curve estimation does not distort well-behaved standard curves and that the recovery is specific to artifact-affected targets. For this clean control the threshold-based efficiency was in fact marginally closer to 100% than the Fit-based value, underscoring that neither estimator is universally preferable. Overall, these data indicate that Fit Cq may offer advantages when evaluating standard curves for amplification efficiency and linearity when threshold miscalls occur.

**Figure 5.**
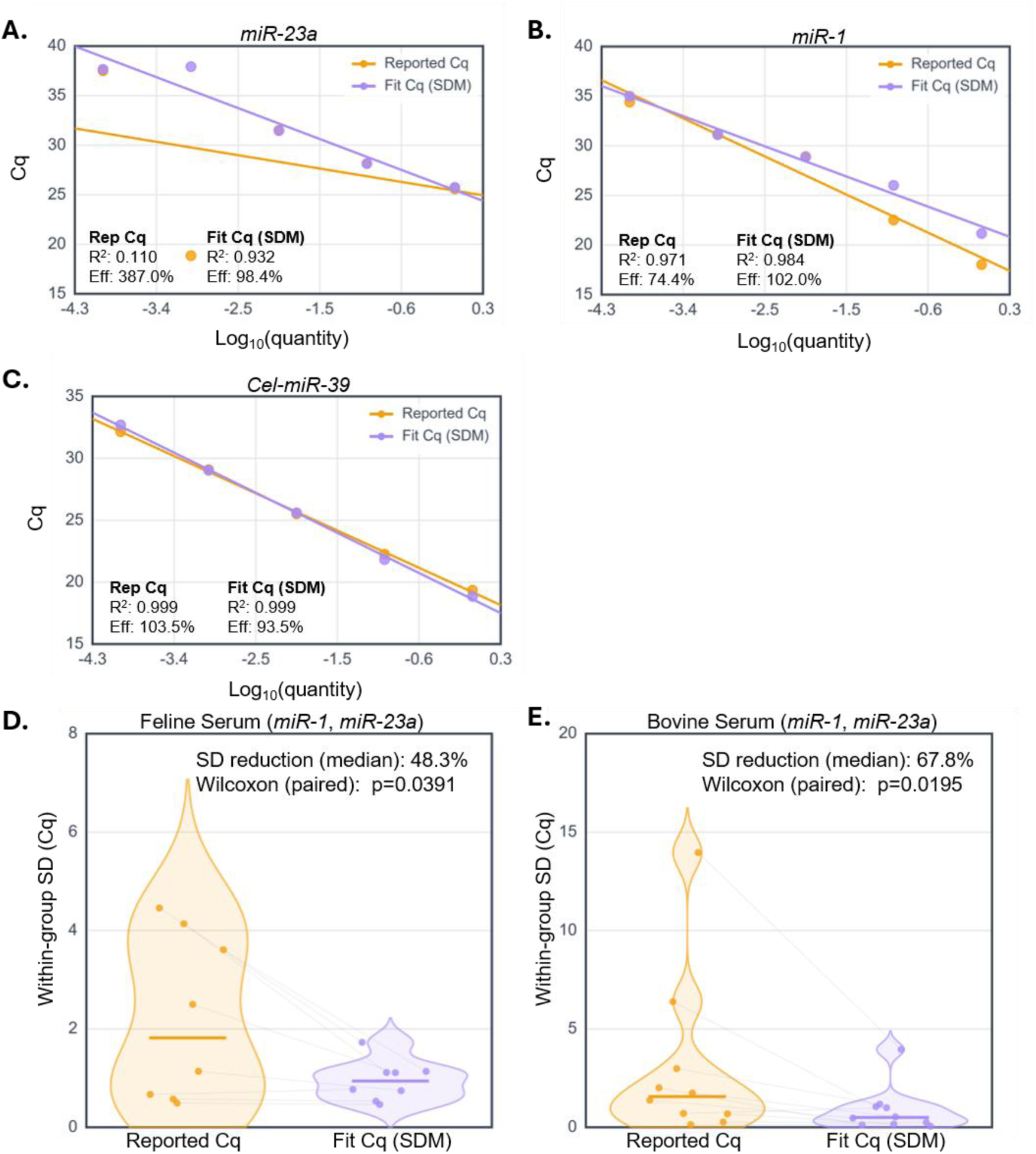
Comparison of amplification efficiency and precision in serum microRNA between Cq calling methods. (A–C) Standard-curve dilution series on pooled feline serum (n=6) from 1 extraction (Cq versus log₁₀ template quantity) for (A) *miR-23a*, (B) *miR-1*, and (C) the co-diluted *cel-miR-39* control, with linear fits, R², and amplification efficiency (Eff) computed separately from Reported Cq (orange) and Fit Cq (SDM; purple). A biphasic miscall in the *miR-23a* series inverts the threshold-based standard curve (efficiency 387%, R² = 0.11), which the Fit Cq restores (98.4%, R² = 0.93); the clean *cel-miR-39* control passes by both methods. (D, E) Bean plots from within-group SD of Cq by method for (D) pooled feline serum of 6 animals (*miR-1* and *miR-23a*; n=4 (independent extractions), run in technical triplicate) and (E) pooled bovine serum (*miR-1* and *miR-23a*; n=5 (independent extractions), run in technical duplicate); horizontal bars, medians; connecting lines link matched groups; median SD reduction and paired Wilcoxon signed-rank p-values are indicated. Eff, amplification efficiency [E = 10^(−1/slope) − 1].

**Table 2.** Well-level QC across a five-point standard-curve dilution series. Reported and Fit Cq (SDM) for *miR-1*, *miR-23a*, and the co-diluted *cel-miR-39* control. Highlighted wells are biphasic threshold miscalls: A1 (*miR-1*, undiluted) and D5 (*miR-23a*, 10⁻³). D5 is the critical case — its threshold value (18.85) reads earlier than the undiluted A5 well (25.58), inverting the dilution series and producing a nonsensical threshold efficiency; Fit Cq (37.92) restores monotonic spacing. *cel-miR-39* is the clean control, concordant by both methods across all dilutions. ^¹^Curve shape determined by five-parameter logistic (5PL) sigmoid fit (Spiess *et al.* 2008; Ritz & Spiess 2008).^²^Reported Cq: the instrument threshold-crossing value as imported; unmodified by qPCR Guru.^³^Fit Cq (SDM): the second-derivative maximum (cpD2) of the fitted 5PL curve (Ritz & Spiess 2008).

| Well | Dilution | Target | Curve Shape | R' (5PL) | Reported Cq | Fit Cq (SDM) | $\Delta Cq$ | Flags |
| --- | --- | --- | --- | --- | --- | --- | --- | --- |
| A1 | 1 | *miR-1* | Abnormal | 0.997 | 18.02 | 21.16 | +3.14 | Biphasic amplification |
| B1 | 0.1 | *miR-1* | Normal | 0.996 | 22.51 | 26.01 | +3.50 | — |
| C1 | 0.01 | *miR-1* | Normal | 0.986 | 28.92 | 28.86 | -0.06 | — |
| D1 | 0.001 | *miR-1* | Normal | 0.984 | 31.16 | 31.11 | -0.05 | — |
| E1 | 0.0001 | *miR-1* | Normal | 0.963 | 34.40 | 34.99 | +0.59 | — |
| A5 | 1 | *miR-23a* | Normal | 1.000 | 25.58 | 25.74 | +0.16 | — |
| B5 | 0.1 | *miR-23a* | Normal | 0.999 | 28.14 | 28.14 | +0.00 | — |
| C5 | 0.01 | *miR-23a* | Normal | 0.997 | 31.50 | 31.46 | -0.04 | — |

| Well | Dilution | Target | Curve Shape' | R' (5PL) | Reported Cq' | Fit Cq (SDM)' | $\Delta Cq$ | Flags |
| --- | --- | --- | --- | --- | --- | --- | --- | --- |
| D5 | 0.001 | *miR-23a* | Abnormal | 0.966 | 18.85 | 37.92 | +19.07 | Biphasic amplification |
| E5 | 0.0001 | *miR-23a* | Normal | 0.973 | 37.50 | 37.65 | +0.15 | — |
| A12 | 1 | *cel-miR-39* | Normal | 1.000 | 19.35 | 18.85 | -0.50 | — |
| B12 | 0.1 | *cel-miR-39* | Normal | 1.000 | 22.26 | 21.82 | -0.44 | — |
| C12 | 0.01 | *cel-miR-39* | Normal | 1.000 | 25.51 | 25.60 | +0.09 | — |
| D12 | 0.001 | *cel-miR-39* | Normal | 1.000 | 29.08 | 29.03 | -0.05 | — |
| E12 | 0.0001 | *cel-miR-39* | Normal | 1.000 | 32.14 | 32.69 | +0.55 | — |

### Evaluation of precision between Cq estimation methods

Finally, we compared within-group precision in pooled serum miRNA samples. Fit Cq reduced the median within-group standard deviation by 48.3% in feline serum (*miR-1* and *miR-23a*; Wilcoxon signed-rank p = 0.039; Figure 5D) and by 67.8% in bovine serum (p = 0.020; Figure 5E). In contrast to the clean Akt data, where precision was comparable between methods, the low-input serum miRNA studies showed a measurable precision advantage for the Fit Cq.

### Interpretation and limitations

Taken together, these results indicate that the two estimators agree on well-behaved, high-expressing targets but diverge where baseline artifacts and biphasic amplification cause the threshold to be crossed before the true exponential rise, conditions that are common in low-input serum miRNA. In these datasets the Fit Cq recovered replicate-concordant values, restored MIQE-compliant amplification efficiencies, and improved precision. Second-derivative-maximum estimation is a well-established alternative to threshold calling [9, 10], and qPCR Guru computes both for every well so that users can compare them easily through direct uploads of instrument exported files. Our data suggest, but do not prove, that the full-curve estimate is the more reliable choice when threshold miscalls are present. Several limitations bound these conclusions. The replicate numbers are small, the precision comparison rests on technical replication across only two serum sources, and the per-curve miscalls are illustrated with individual wells rather than large cohorts. The efficiency and precision benefits should therefore be read as evidence that the estimator choice can matter materially for low-input miRNA, not as a general claim that full-curve estimation is superior under all conditions. On clean targets the two approaches were equivalent, and for one clean control the threshold estimate was marginally better.

## CONCLUSIONS

We describe qPCR Guru, a free browser-based platform that enables comparison between threshold-based and full-curve quantification cycle estimates for every well. On well-behaved, high-expressing targets the two estimates were concordant and produced equivalent relative quantification, indicating that full-curve estimation can be applied without distorting clean data. In low-expressing serum microRNA, where baseline artifacts and biphasic amplification caused threshold miscalls, the full-curve estimate recovered replicate-concordant values, restored interpretable amplification efficiencies, and improved precision. Because both estimates are reported together, users can identify unreliable calls and select the appropriate value rather than committing to a single method in advance. These findings, although based on a limited number of replicates and sample sources, indicate that the choice of cycle estimator can meaningfully affect quantitative results for low-input microRNA.

## FUTURE PERSPECTIVE

As microRNA profiling in biofluids continues to expand in biomarker research and clinical diagnostics, the reliability of low-input quantification will become increasingly consequential. Routine, side-by-side comparison of threshold and full-curve cycle estimates, together with automated flagging of the disagreements between them, could help standardize the detection of unreliable calls that currently pass unnoticed. We anticipate that accessible, browser-based analysis tools will lower the practical barriers to such curve-level scrutiny and to MIQE-aligned reporting more broadly. Larger studies incorporating biological replication across additional biofluids and species will be needed to define the conditions under which full-curve estimation offers the greatest benefit, and to establish whether its precision and efficiency advantages translate into more reproducible downstream biological conclusions.

---

**Supplementary Figure 1.**
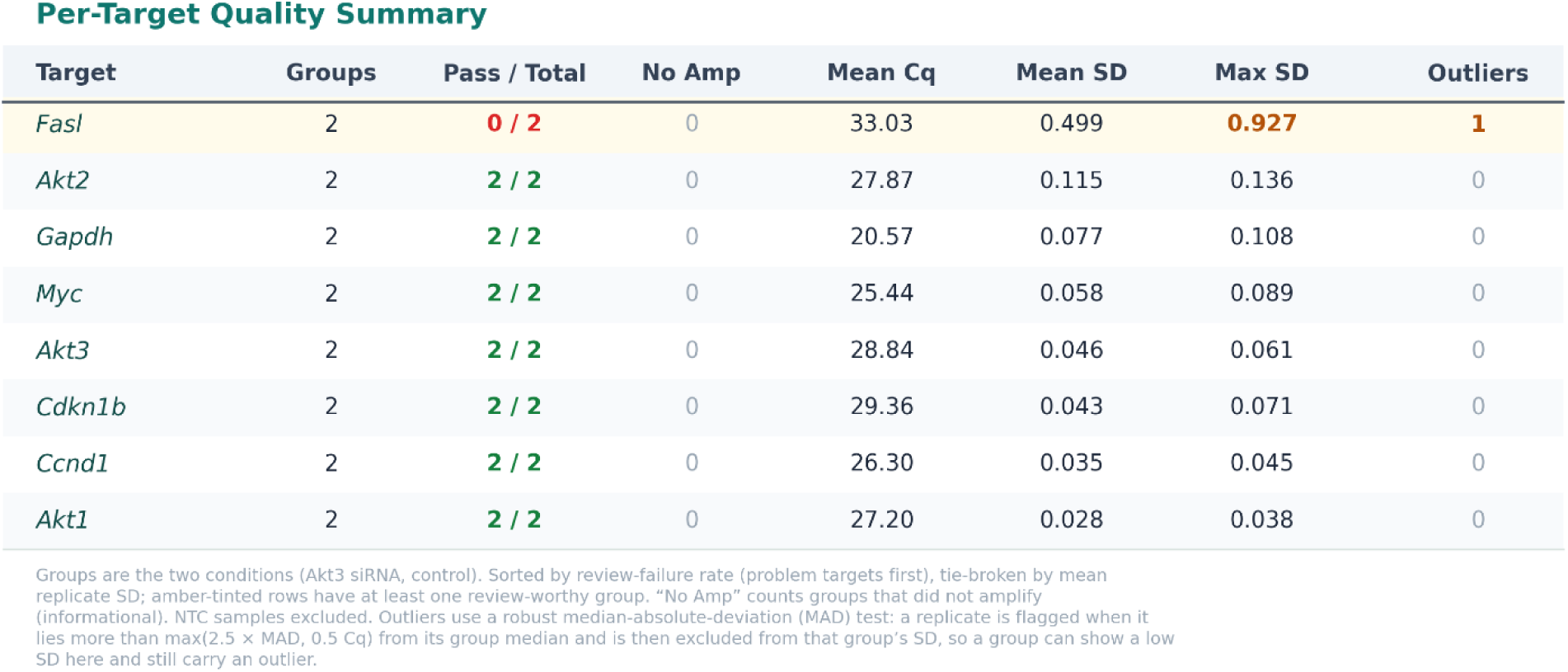
Per-target data-quality summary for the Akt3 knockdown experiment. For each target, the platform summarizes quality across the two sample groups (Akt3 siRNA and control): the number of groups detected, groups passing quality review (Pass / Total), groups that did not amplify (No Amp), the mean Cq across groups, the mean and maximum within-group replicate standard deviation (Mean SD, Max SD; in cycles), and the number of replicate outliers. Rows are sorted by review-failure rate (problem targets first). Outliers are identified by a robust median-absolute-deviation (MAD) test (a replicate flagged when it lies more than max(2.5 × MAD, 0.5 Cq) from its group median) and are excluded from that group’s standard deviation; consequently the *Fasl* control standard deviation reflects two replicates after exclusion of one flagged outlier (Cq 34.66). *Fasl* was the only target with a review-worthy group, driven by elevated replicate variability and the excluded control outlier. Cq, quantification cycle; SD, standard deviation; MAD, median absolute deviation; NTC, no-template control (excluded).

**Supplementary Figure 2.**
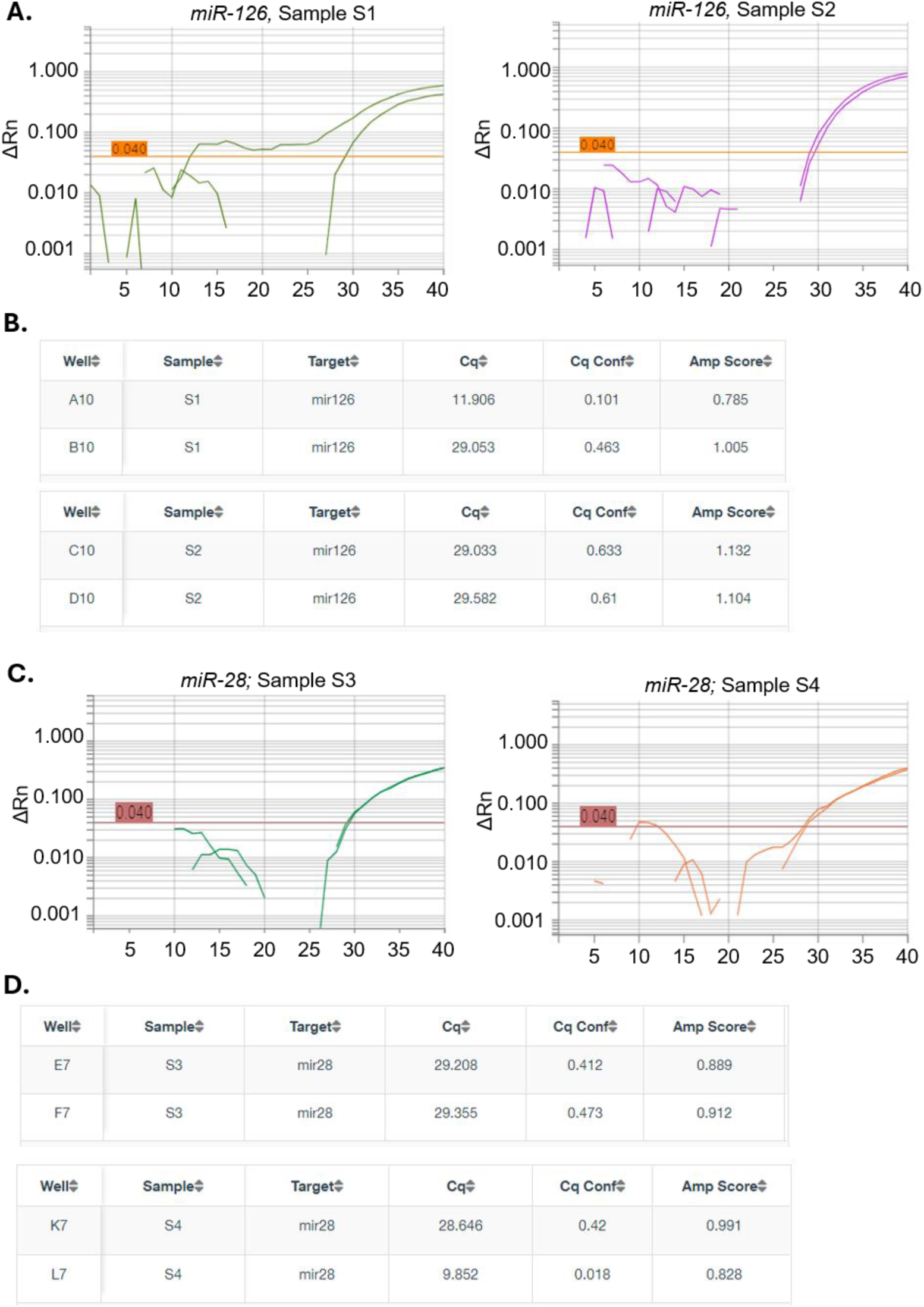
Instrument amplification plots and amplification-confidence scores for the serum microRNA miscall wells. (A, C) QuantStudio ΔRn-versus-cycle plots for the *miR-126* (S1, S2) and *miR-28* (S3, S4) wells from individual feline serum samples. (B, D) Corresponding instrument Cq, amplification-confidence (Cq Conf), and amplification (Amp Score) values; the miscalled wells A10 and L7 show low amplification confidence (0.101 and 0.018) relative to their normal technical replicates (∼0.42-0.46).

**Supplementary Table 1. Quantstudio Export of the Akt3 knockdown in CT-2A mouse glioma cells.**

**Supplementary Table 2. qPCR Guru computed ddCq results from either Reported Cq or Fit Cq.**

**Supplementary Table 3. Quantstudio Export from the individual feline serum miscalls of *miR-126* and *miR-28*.**

**Supplementary Table 4. Quantstudio Export from the pooled feline serum dilution series.**

**Supplementary Table 5. Quantstudio Export from the pooled feline serum replication analysis**

**Supplementary Table 6. Quantstudio Export from the pooled bovine serum replication analysis.**

## DISCLOSURES

### Competing interests

A.M.S. and O.S. are founders and owners of Amos Innovations, LLC, the developer of qPCR Guru; the platform is free to use. R.C. is a co-founder of MI:RNA Diagnostics, Ltd. S.L.S. is a founder and owner of Aruna Bio, Inc. M.M.S. declares no competing interests.

### Funding

This work received no grant funding.

### AI / writing-assistance disclosure

During the development of qPCR Guru and the preparation of this manuscript, the authors used Claude (Anthropic), Claude Opus 4.6, 4.7, and 4.8 for software development; Claude Opus 4.8 for manuscript language editing. All AI-assisted outputs, including software code and manuscript text, were reviewed, tested, and edited by the authors, who take full responsibility for the content. Figures were generated programmatically from the experimental data and are not AI-generated images.

### AUTHOR CONTRIBUTIONS (CRediT)

A.M.S.: supervision, conceptualization, methodology, software, formal analysis, investigation, writing — original draft. O.S.: conceptualization, software, methodology. M.M.S.: investigation, methodology. R.C.: supervision, resources. S.L.S.: supervision, resources. All authors: writing — review & editing.

## ACKNOWLEDGEMENTS

The authors thank all members of the MI:RNA team and Dr. Stice’s laboratory members for their support.

## DATA AVAILABILITY STATEMENT

Raw instrument exports are provided as supplementary tables. The qPCR Guru application (qpcrguru.com) is available for free use.

## Notes

### Competing Interest Statement

The authors have declared no competing interest.

